# Cortical dynamics of speech feedback control in non-fluent Primary Progressive Aphasia

**DOI:** 10.1101/2022.07.28.501928

**Authors:** Hardik Kothare, Kamalini G. Ranasinghe, Leighton B. Hinkley, Danielle Mizuiri, Abigail Licata, Michael Lauricella, Susanne Honma, Valentina Borghesani, Corby Dale, Wendy Shwe, Ariane Welch, Zachary Miller, Maria Luisa Gorno-Tempini, John F. Houde, Srikantan S. Nagarajan

**Author notes:** Correspondence to: Srikantan Nagarajan, PhD, 513 Parnassus Ave, S679, San Francisco, CA 94143, USA.

## Abstract

Primary Progressive Aphasia (PPA) is a clinical syndrome in which patients progressively lose speech and language abilities. The non-fluent variant of PPA (nfvPPA) is characterised by impaired motor speech and agrammatism. To date, no study in nfvPPA patients has either examined speech motor control behaviour or imaged the speech motor control network during vocal production. Here, we did this using a novel structure-function imaging approach integrating magnetoencephalographic imaging of neural oscillations with voxel-based morphometry (VBM). We examined task-induced non-phase-locked neural oscillatory activity during a vocal motor control task, where participants were prompted to phonate the vowel /□/ for ∼2.4s while the pitch of their auditory feedback was shifted either up or down by 100 cents for a period of 400ms mid-utterance. Participants were 18 nfvPPA patients (14 female, mean age = 67.79 ± 8.02 years) and 17 controls (13 female, mean age = 64.81 ± 5.76 years). Patients showed a smaller compensation response to pitch perturbation than controls (p < 0.05). Task-induced neural oscillations across five frequency bands were reconstructed in source space for each subject during pitch feedback perturbation. Patients exhibited reduced task-induced alpha-band (8-12Hz) neural activity unrelated to their atrophy patterns, in the right temporal lobe and the right temporoparietal junction (p < 0.01) from 250ms to 750ms after pitch perturbation onset. Patients also showed increased task-induced beta-band (12-30Hz) activity also unrelated to cortical atrophy in the left dorsal sensorimotor cortex, left premotor cortex and the left supplementary motor area (p < 0.01) from 50ms to 150ms after pitch perturbation onset. Reduced average alpha-band power at the peak voxel in the temporoparietal cluster in the right hemisphere could predict speech motor impairment in patients (*β* = 3.41, F = 8.31, p = 0.0128) whereas increased average beta-band power at the peak voxel in the left dorsal sensorimotor cluster could not (*β* = -1.75, F = 1.72, p = 0.2123). Collectively, these results suggest significant disruption in sensorimotor integration during vocal production in nfvPPA patients which occurs unrelated to patterns of atrophy. These findings highlight how multimodal structure-function imaging in PPA enhances our understanding of its pathophysiological sequelae.

## Introduction

Primary progressive aphasia (PPA) is a clinical syndrome characterised by progressive loss of speech and language abilities.^1^ Currently accepted clinical classification identifies three subtypes of PPA,^2^ each with a characteristic impairment in speech and language function, namely: a logopenic variant of PPA (lvPPA), a semantic variant of PPA (svPPA) and a non-fluent variant of PPA (nfvPPA). Among these three variants, the nfvPPA subtype includes patients who predominantly exhibit motor-speech deficits including apraxia and agrammatism. As a neurodegenerative disorder, patients with nfvPPA exhibit signature patterns of neuronal atrophy in structural magnetic resonance imaging (MRI) along the left posterior fronto-insular region - encompassing areas involved in the production of speech and language like the inferior frontal gyrus (IFG), insula, premotor cortex and supplementary motor area (SMA).^3-5^ Non-invasive neuroimaging techniques have also shown in nfvPPA significant alterations in functional and structural connectivity between regions of the speech production network.^6-8^ Both behavioural and neuroanatomical distinctions in nfvPPA are the most prominent in the later stages of the disease, making diagnosis challenging in the early stages of nfvPPA. Ultimately, how these observed deficits in activation and connectivity are influenced by neurodegeneration and directly translate into speech motor impairments is poorly understood.

One important function of the speech system is sensorimotor control, where an individual will respond to changes in sensory feedback during ongoing speaking. Current models of speech production posit that during speaking, sensory feedback is compared with a motor-derived prediction of that feedback, with any mismatches driving compensatory adjustments to speech motor output.^9, 10^ Under normal speaking conditions, sensory feedback usually matches predictions, resulting in minimal feedback prediction errors and minimal compensatory responses. However, if the vocal tract is perturbed from its normal state (by, for example, muscle fatigue or food in the mouth), incoming sensory feedback will mismatch predictions, resulting in feedback prediction errors being detected by the speech sensorimotor control system, which usually responds by generating a compensatory motor adjustment to ongoing speaking. This process of detecting and compensating for feedback prediction errors during speaking can also be seen in the laboratory when sensory feedback is externally perturbed. A well-known example of this occurs when the pitch of a speaker’s auditory feedback is artificially perturbed during speaking, causing most speakers to shift their produced pitch in a direction opposite that of the perturbation. This compensatory response is involuntary and is called the pitch perturbation reflex.^11^ Its psychophysical characteristics and neural basis have been widely studied both in healthy controls^12-16^ and in a number of neurological disorders.^17-20^ making it an ideal experimental paradigm for studying the dynamics of sensorimotor integration in patients with nfvPPA.

Using a novel structure-function imaging approach, we examined here the behavioural response during pitch perturbation reflex and underlying neurophysiological characteristics of sensorimotor integration in patients with nfvPPA. We aimed to test the hypothesis that alterations in the pitch perturbation reflex cannot be explained by neurodegeneration of sensorimotor control regions alone. Specifically, we compared vocal and neural responses induced by perturbing the pitch of auditory feedback in patients with nfvPPA against the same processes in healthy controls. We employed here a novel multimodal structure-function approach, which leveraged source reconstructions with a high temporal resolution in magnetoencephalography (MEG) with voxelwise measures of cortical atrophy (as a covariate) obtained using volumetric MRI. We predict that given the pathophysiological processes in nfvPPA that target brain regions involved in speech production, nfvPPA patients will exhibit differences in behavioural and neuronal oscillations during the pitch perturbation reflex that cannot be explained by reductions in cortical atrophy alone. We further predicted that the magnitude of this pitch response would be associated with severity of speech motor impairment in nfvPPA patients as measured by standard clinical neuropsychological assessments.

## Materials and methods

### Participants

For this study, 20 patients and 18 healthy controls (see Table 1 for demographic details) were recruited from the Memory and Aging Center at the University of California, San Francisco (UCSF). 17 patients and 14 controls took part in the pitch perturbation experiment with simultaneous MEG imaging. Three patients and four controls took part in the behavioural pitch perturbation experiment without neuroimaging. Patients’ PPA diagnosis and classification as nfvPPA was conducted by a team of expert neurologists. Eligibility criteria for controls included: no structural brain abnormalities, normal cognitive performance, absence of neurological or psychiatric disorders. Participants (or their assigned surrogate decision makers) provided informed consent before taking part in the study. This study was approved by the Committee on Human Research of the University of California, San Francisco.

**Table 1.**
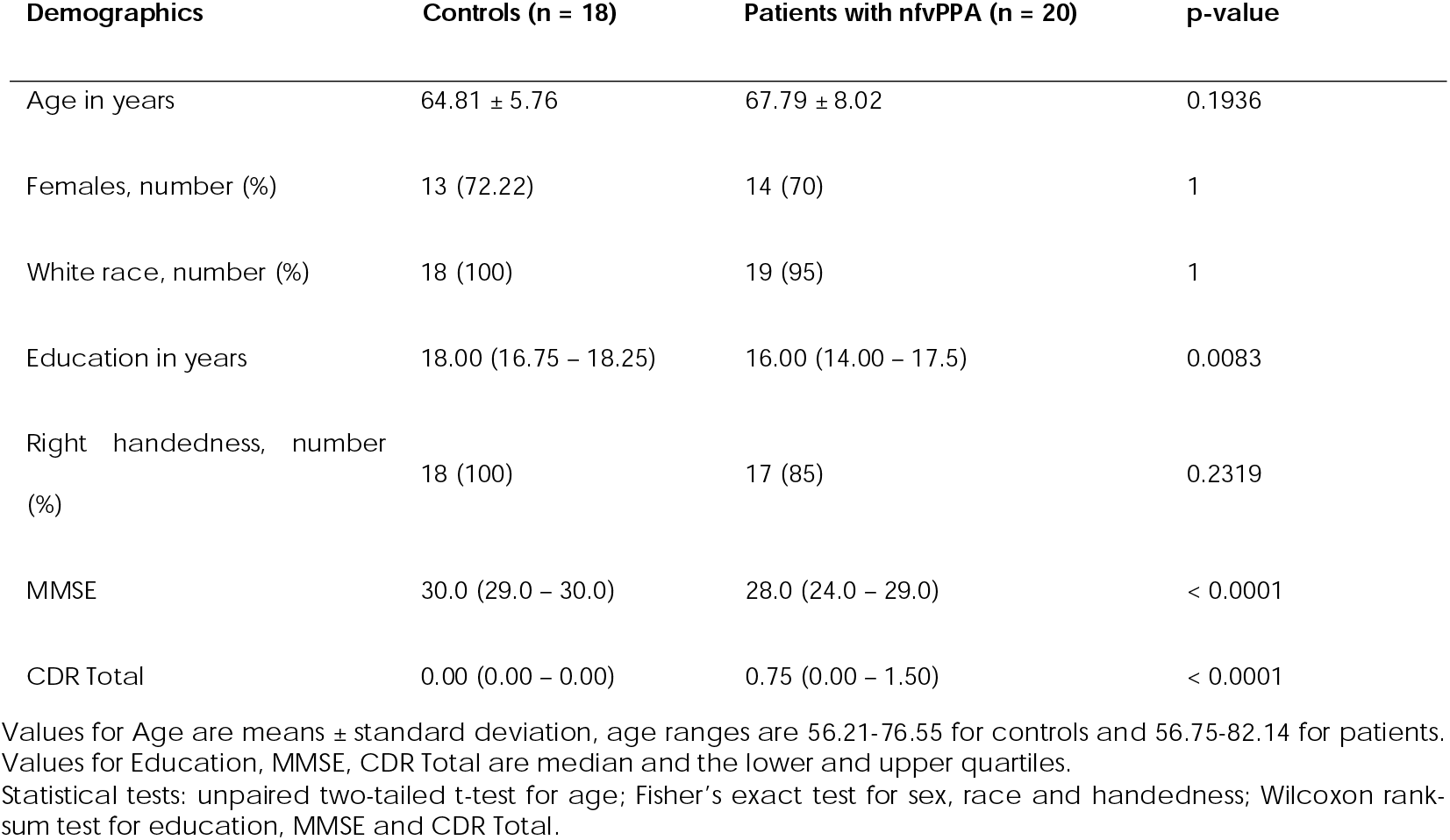
Participant demographics.

| Demographics | Controls (n = 18) | Patients with nfvPPA (n = 20) | p-value |
| --- | --- | --- | --- |
| Age in years | 64.81 ± 5.76 | 67.79 ± 8.02 | 0.1936 |
| Females, number (%) | 13 (72.22) | 14 (70) | 1 |
| White race, number (%) | 18 (100) | 19 (95) | 1 |
| Education in years | 18.00 (16.75 – 18.25) | 16.00 (14.00 – 17.5) | 0.0083 |
| Right handedness, number (%) | 18 (100) | 17 (85) | 0.2319 |
| MMSE | 30.0 (29.0 – 30.0) | 28.0 (24.0 – 29.0) | < 0.0001 |
| CDR Total | 0.00 (0.00 – 0.00) | 0.75 (0.00 – 1.50) | < 0.0001 |

Values for Age are means ± standard deviation, age ranges are 56.21-76.55 for controls and 56.75-82.14 for patients. Values for Education, MMSE, CDR Total are median and the lower and upper quartiles. Statistical tests: unpaired two-tailed t-test for age; Fisher's exact test for sex, race and handedness; Wilcoxon rank-sum test for education, MMSE and CDR Total.

### Neuropsychological assessment

All participants underwent a Mini-Mental State Examination (MMSE). A Clinical Dementia Rating (CDR) score was also calculated for all participants after an interview with the participants and their caregivers. ^21^ 19 of the 20 patients also underwent a comprehensive neuropsychological evaluation to test their performance on a number of speech and language tasks as described elsewhere^22, 23^ (Supplementary Table 1). The Western Aphasia Battery was used to evaluate speech and syntactic production.^24^ Apraxia of speech and dysarthria were rated using Motor Speech Evaluation.^25^ To measure long and short syntax comprehension, sentences were read aloud by the examiner and the patient had to select the picture that best matched the sentence from two options. Confrontation naming was assessed using a short form version (15 items) of the Boston Naming Test.^26^ Patients were asked to name as many animals as possible in 60 seconds to assess category fluency.^27^ A subset of 16 items from the Peabody Picture Vocabulary Test (PPVT; 4 items each: verbs, descriptive, animate and inanimate) where patients were asked to match a word with one of four picture choices was used to test word comprehension.^28^

### Structural MRI acquisition

Participants who underwent an MEG scan also had a structural MRI of their brains acquired. The MRI acquisition took place in a 3T Siemens MRI scanner at UCSF. T1-weighted structural MRI images were acquired using a T1-MPRAGE sequence (repetition time = 2300ms, echo time = 2.98ms, inversion time = 900ms, slice thickness = 1mm, field of view = 240mm x 256mm). These structural MRIs were used to generate head models for source-space reconstruction of MEG sensor data and to calculate grey matter volume estimates for statistical correction of grey-matter atrophy in the MEG results. Individual structural MRIs were spatially normalised to a standard Montreal Neurological Institute (MNI) template for visualisation and cohort-level statistical analyses using SPM8.

### MEG Imaging

A 275-channel whole-head MEG scanner (CTF Inc., Coquitlam, BC, Canada) was used for MEG imaging. Participants were scanned in supine position. Signals were acquired at a sampling rate of 1200Hz and three fiducial coils (one at the nasion and two at preauricular points on both sides) were used to record head position relative to the sensor array. These fiducial locations were co-registered to participants’ structural MRI images and head shapes were generated for each individual.

### Pitch perturbation experiment

The pitch perturbation experiment consisted of 120 trials. In every trial, participants were prompted to vocalise the vowel /□/ for as long as they saw a green dot on the screen (duration ∼ 2.4s) (see Fig. 1A). Participants’ vocal output was recorded using an MEG-compatible optical microphone (Phone-Or Ltd., Or-Yehuda, Israel) and they could simultaneously hear themselves through MEG-compatible insert earphones (ER-3A, Etymotic Research, Inc., Elk Grove Village, IL). During every trial, between 200-500ms after voice onset was detected, the pitch of the participants’ auditory feedback was perturbed either upwards or downwards by 100 cents (1/12^th^ of an octave) for a duration of 400ms using a real-time digital signal processing program called Feedback Utility for Speech Production (FUSP), which runs in Linux on a PC and has also been used in previous studies.^19, 29-31^ The direction of shift was randomly determined for each trial with an equal number of trials with upward and downward shifts.

**Figure 1:**
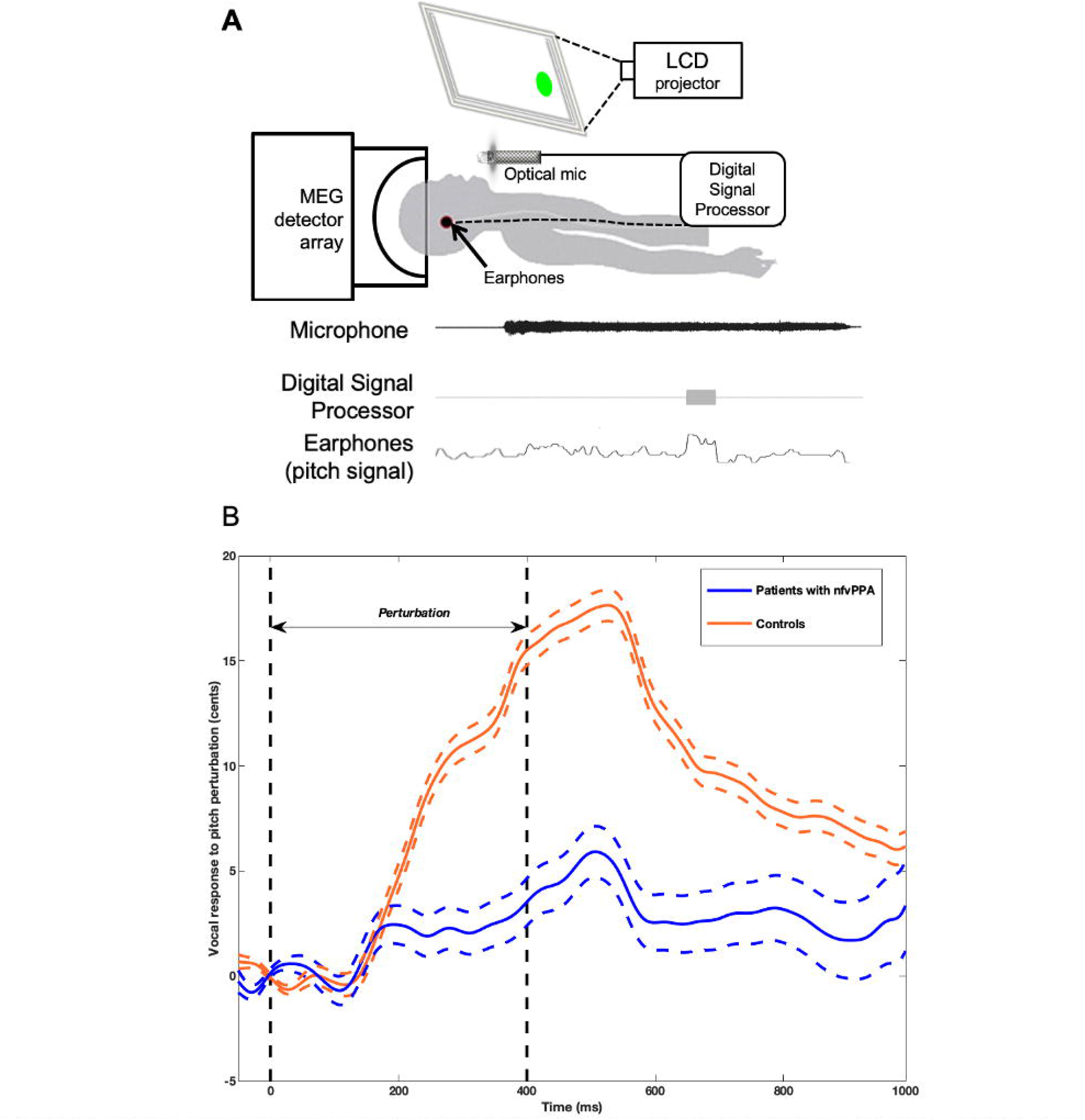
Task description and behavioural results. **(A)** During every trial, participants started to vocalise the vowel /□/ into a microphone upon seeing a green dot on the display. Participants also wore earphones through which they could hear themselves throughout the trial. Between 200-500ms after the start of vocalisation, a digital signal processing unit shifted the pitch of the participants’ speech signal either up or down for 400ms and sent this shifted signal to the participants’ earphones. **(B)** The x-axis depicts time from perturbation onset (ms) and the y-axis depicts vocal pitch (cents). Pitch feedback perturbation lasted for 400ms. On average, patients with nfvPPA (n = 18) have a smaller compensation response to pitch perturbation as compared to controls (n = 17). Dashed lines represent standard error of the mean.

### Data Analysis

#### Audio Data Processing

Participants’ speech and the pitch-altered feedback were both recorded at 11,025 Hz. For every trial, the time course of the pitch of the participant’s speech (the pitch track) was determined using an autocorrelation-based pitch estimation method.^32^ Pitch tracks for each trial were aligned to perturbation onset, and ranged from 200ms prior to perturbation onset to 1000ms after perturbation onset. All trials with pitch tracking errors and incomplete utterances were marked bad and excluded from further analysis. For all good trials, pitch values were converted from Hertz to cents using the following formula:

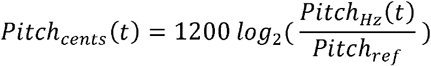

where *Pitch*_*Hz*_(*t*) is the pitch value in Hertz at timepoint t and *Pitch*_*ref*_ is the reference pitch calculated as the mean pitch over a window spanning from 50ms prior to perturbation onset to 50ms after perturbation onset. Participants’ responses to pitch perturbation were expressed as deviations from their baseline pitch track. For each participant, responses to both upward and downward perturbations were calculated and pooled together. Downward responses to upward perturbations were flipped and combined with upward responses to downward perturbations, thus making all compensatory responses positive.

For statistical analysis of vocal pitch data, the pitch track for each individual trial was divided into bins of 50ms each and pitch values were averaged within these bins. Group differences in compensatory pitch responses between patients and controls were calculated for each of these bins using trial means from both groups and running a one-way analysis of variance (anova1 function, MATLAB, MathWorks, Natick, MA). To control for Type I error, Bonferroni thresholds were applied to p-values for α = 0.05.

#### MEG Data Processing

We performed third-order gradient noise and direct current (DC) offset correction on the MEG sensor data. Pitch perturbation onset was marked for every trial and subsequent neural analyses were locked to this perturbation onset marker. All trials were visually inspected; trials with noisy signal (>1pT) due to head movement, dental artefact, eye blinks or saccades were excluded from the analysis. We examined task-induced non-phase-locked neural oscillatory responses in the theta (4-7Hz), alpha (8-12Hz), beta (13-30Hz), low gamma (30-55Hz) and high gamma (65-150Hz) frequency bands. However, we found cohort differences only in alpha and beta bands which are known to contain robust electrophysiological signatures of sensorimotor integration and control.^33, 34^ Hence, we focus on these two bands in the results and the discussion.

Source localisation for induced alpha-band and beta-band activity in each participant was performed using time-frequency-optimised adaptive spatial filtering (8mm lead field) in the Neurodynamic Utility Toolbox for MEG (NUTMEG: https://www.nitrc.org/projects/nutmeg).^35, 36^ Voxel-by-voxel estimates of neural activity were generated using a linear combination of a spatial weight matrix and a sensor data matrix. Active windows were defined as the time period after perturbation onset. Pseudo-F statistics were computed by comparing the active window to a control ‘baseline’ window prior to perturbation onset (alpha band: window length = 400ms, step size = 100ms, beta band: window length = 200ms, step size = 50ms). Using the NUTMEG toolbox,^37^ both within-group and between-group statistical analyses were performed using statistical non-parametric mapping methods. For within-group contrasts, we used a 5% False Discovery Rate (FDR) to correct for multiple comparisons across time and space and corrected p-value thresholds were calculated for α = 0.01. To further control for false positive activations, cluster correction was also performed to exclude clusters with less than 40 contiguous voxels.

When comparing differences in oscillatory power between groups (nfvPPA vs. HC) corrected for atrophy, we used the Nutmeg Atrophy Statistics (NAS) toolbox in NUTMEG. Briefly, NAS takes voxelwise values of grey matter atrophy as a covariate when comparing across groups. Grey matter (GM) maps were generated by tissue-type segmentation of the T1-weighted MRIs of patients and controls using the DARTEL pipeline^38^ and spatially normalised to a custom group template (n = 100) in MNI space using the same dimensions as the MEG data (79×95×68 matrix). With this information aligned at the voxel level, differences between groups in oscillatory (i.e. alpha, beta) power were estimated using a voxelwise analysis of covariance (ANCOVA) with GM values as a covariate. We used the same FDR correction parameters as in the within-group analyses (p<0.01), but to allow for increased sensitivity for cohort comparisons, the threshold used for subsequent cluster correction was 20 voxels.

#### Correlation of average peak neural activity with speech motor impairment

To explore possible relationships between neural responses to pitch perturbation and speech motor impairment in patients, we calculated a Speech Motor Composite Score (SMCS), for the 16 patients having both neural data and neuropsychological assessment scores, using the following formula:

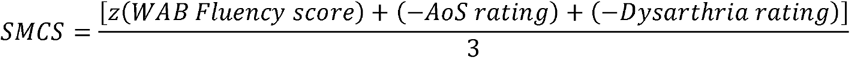

where z stands for z-score normalisation with respect to the normative score, WAB = Western Aphasia Battery, AoS = Apraxia of Speech. The values for AoS rating and Dysarthria rating were negated because we wanted lower scores to depict greater impairment and in both these ratings higher values indicate greater impairment. AoS and Dysarthria ratings are already normalised and hence a z-score was not calculated for them.

For the voxel and timepoints showing peak significant differences between patients and controls, the average alpha power and the average beta power were calculated for each of the 16 patients. Then, for each frequency band (alpha and beta), a generalised linear model (GLM) was fit in SAS 9.4 (SAS Institute Inc., Cary, NC) to investigate whether the average peak power in the frequency band predicted speech motor impairment.

### Data availability

The data that supports the findings of this study will be made available upon reasonable request.

## Results

### Participant characteristics

Patients with nfvPPA showed mild cognitive impairment (*median value for MMSE* = 28, see Table 1). The control and patient cohorts did not differ in terms of age, sex, handedness and race (see Table 1). However, controls and patients did differ when it came to average years of education (*median* = 18 for controls and 16 for patients).

### Behavioural response to pitch perturbation

Figure 1B shows both groups’ vocal response to pitch perturbation. Patients with nfvPPA and controls started responding to perturbation at ∼150 ms and followed a similar trajectory until ∼200 ms. From 200ms onwards, control participants continued to respond along the same incline until they reached a peak at 526ms (*peak response* =17.64 ± 0.75 cents). Patients with nfvPPA, in sharp contrast, showed a different trajectory after ∼200ms where they followed a less-steep slope and reached a smaller peak at 505ms (*peak response* = 5.91 ±1.22 cents). From 200ms until 950ms after perturbation onset, patients with nfvPPA showed a significantly reduced behavioural response compared to controls (Figure 1B).

To evaluate whether the smaller responses in patients was due to a limited vocal motor range, we quantified vocal motor output capacity as pitch variability in patients and controls in a 200ms pre-perturbation baseline window as done in previous studies.^17, 29^ Variability in patients with nfvPPA did not differ significantly from that in controls, for both within-trial and across-trial analyses (Supp. Fig. 1).

### Neural response to pitch perturbation

Next, we examined neural activity patterns during the pitch perturbation response. We found cohort differences only in alpha (8-12 Hz) and beta (13-30 Hz) bands which are known to contain robust electrophysiological signatures of sensorimotor integration.^33, 34^ Hence, we focus on these two bands in the results and the discussion.

As the behavioural response started at ∼150ms (Figure 1B), the neural activity before this point in time can be presumed to be involved in feedback error detection and preparation for motor correction. Neural activity after 150ms until the peak behavioural response is reached (i.e., around 500ms) would reflect processing related to the sensorimotor integration processing of the perturbed auditory feedback – i.e., the continued detection of a 100 cent pitch feedback prediction error (the magnitude of the perturbation) and increasing compensatory motor correction of the pitch of the output speech. Any neural activity after peak compensation would be indicative of the return to baseline vocalisation – i.e., a return to the original intended pitch, during which time there is no longer any feedback prediction error.

### Patients with nfvPPA have impaired right-hemispheric alpha-band activity during speech motor integration processing

Controls showed increased alpha-band activity in bilateral posterior cortices (Figure 2A) throughout the analytical time window of 0-900ms after perturbation onset. This response was spatially larger in the right hemisphere than the left. Regions active during this time window include the bilateral posterior superior parietal cortex and occipital cortex, temporo-parietal junction and the inferior temporal cortices of the right hemisphere. Controls also showed reduced alpha-band power in the left anterior temporal lobe from 250-550ms.

**Figure 2:**
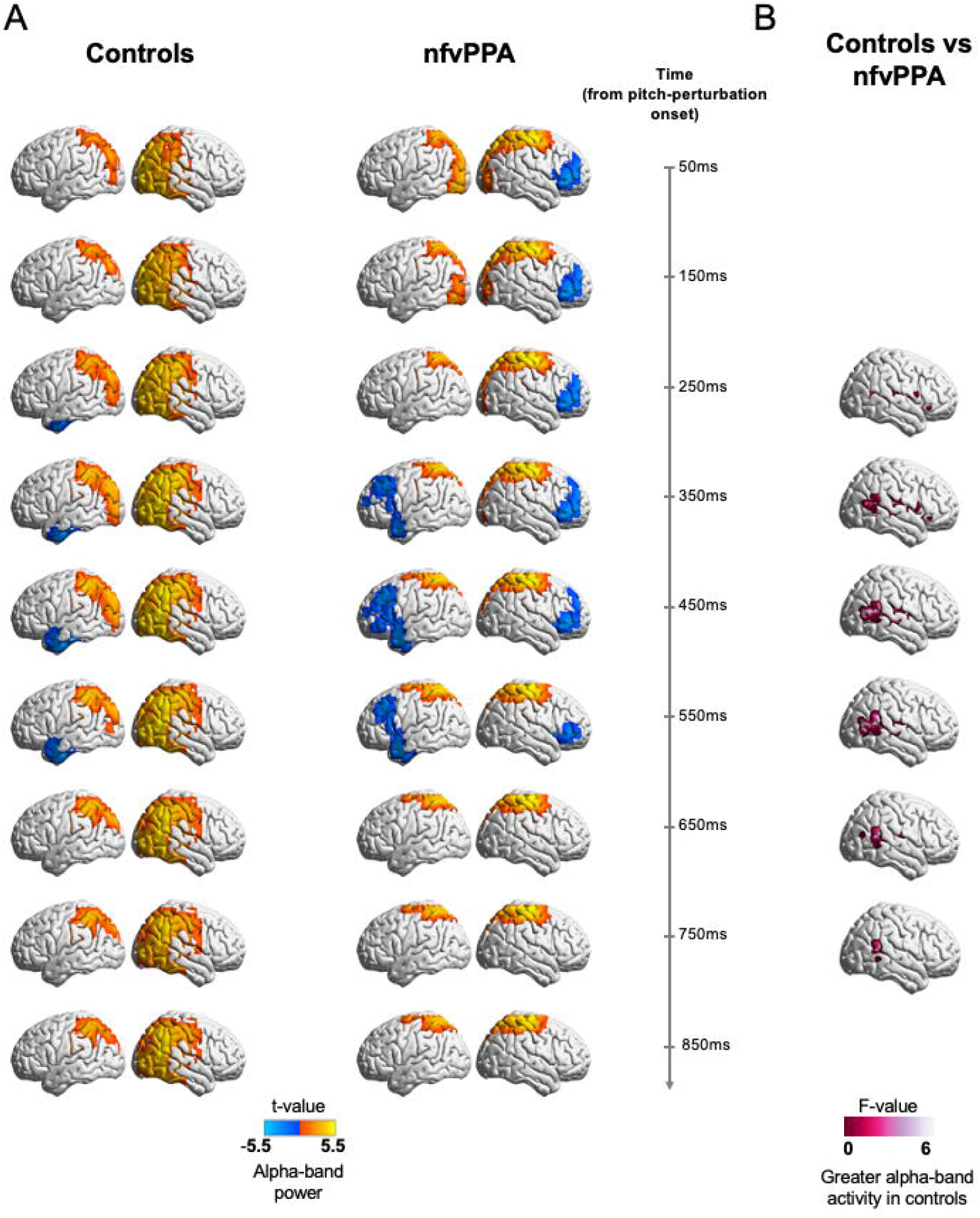
Neural activity during pitch feedback perturbation in alpha band (8 - 12 Hz) **(A)** Neural activity in controls and patients with nfvPPA in the alpha band (8-12 Hz) locked to pitch perturbation onset. **(B)** Patients with nfvPPA (n = 17) show significantly lesser alpha-band activity in the posterior right temporal lobe and the right temporo-parieto-occipital junction as compared to controls (n = 14). Neural activity shown was corrected for grey-matter atrophy and was False Discovery Rate (FDR)-corrected for multiple frequency bands and time points. Cluster correction was also performed at a threshold of 20 voxels and p < 0.01. Timepoint 0 corresponds to perturbation onset.

Patients with nfvPPA also showed increased alpha-band activity in the posterior regions of the brain bilaterally and throughout the analytical window (Figure 2A). However, unlike the controls, a right hemisphere asymmetry was not observed in the nfvPPA patients, although bilateral posterior superior parietal cortices were active. When these time-frequency windows are contrasted between the two groups and corrected for atrophy at the voxel level, right lateralized deficits in nfvPPA are statistically significant (p<0.01 5% FDR), with reduced alpha-band activity in the right temporo-parieto-occipital junction, particularly the right angular gyrus and supramarginal gyrus, in these patients when compared to healthy controls (Figure 2B, Table 2). This difference persisted from 250ms to 750ms after perturbation onset, a window overlapping with neural processing of speech-motor-integration. Although on average, activation in left anterior temporal cortex and frontal cortex (350-550ms) and right inferior frontal cortex (50-550ms) was lower in nfvPPA, this difference was not statistically significant when compared to controls.

**Table 2.** Peak voxels with significant activity differences between patients with nfvPPA and controls.

| Band | Hemisphere | Peak | MNI Coordinates | Anatomical Labels | Time with respect to perturbation |
| --- | --- | --- | --- | --- | --- |
| Alpha (8-12Hz) | Right | 1 | [32 30 -8] | Pars Orbitalis (BA 47) | +250ms |
|  |  | 2 | [50 -31 9] | Superior Temporal Gyrus (BA 22) | +250ms |
|  |  | 3 | [58 10 12] | Operculum (BA 44) | +250ms |
|  |  | 4 | [32 28 -8] | Pars Orbitalis (BA 47) | +350ms |
|  |  | 5 | [57 -21 14] | Supramarginal Gyrus (BA 40) | +350ms |
|  |  | 6 | [50 -49 18] | Angular Gyrus (BA 39) | +350ms |
|  |  | 7 | [48 -47 19] | Angular Gyrus (BA 39) | +450ms |
|  |  | 8 | [53 -18 8] | Primary Auditory region (BA 41) | +450ms |
|  |  | 9 | [53 -21 14] | Supramarginal Gyrus (BA 40) | +450ms |
|  |  | 10 | [50 -47 17] | Angular Gyrus (BA 39) | +550ms |
|  |  | 11 | [50 -47 17] | Angular Gyrus (BA 39) | +650ms |
|  |  | 12 | [52 -47 14] | Angular Gyrus (BA 39) | +750ms |
| Beta (12-30Hz) | Left | 1 | [-28 -13 73] | Premotor Cortex (BA 6) | +50ms |
|  |  | 2 | [-18 -7 76] | Premotor Cortex (BA 6) | +150ms |

BA: Brodmann Area

### Patients with nfvPPA have increased left dorsal sensorimotor beta-band activity during feedback error detection and corrective motoric preparation

Control participants showed increased beta-band activity (Figure 3A) in both hemispheres involving the posterior parietal regions during most of the analytical window (0-700ms post perturbation onset). This increased activity in the parietal cortex in both hemispheres progressed along the primary somatosensory and primary motor cortices from 150 to 550ms. Apart from this robust and consistent activity over the posterior parietal cortices, there was transient early involvement of left frontal regions (50-150 ms), left occipital regions (50-150 ms), right premotor cortex (250-350 ms) and right temporal cortex (350-450 ms). While patients with nfvPPA also showed this pattern of increased beta-band activity over the parietal cortex the spatial distribution of this activity was more dorsally-dominant compared to controls (Figure 3A). The increased beta activity in patients persisted throughout the analytical window in the right hemisphere, while it only persisted from 50-350 in the left hemisphere. At 450 ms, there was a decrease in beta activity in the left inferior frontal cortex in patients but this decrease was not statistically different from that in controls. When beta activity is compared between controls and nfvPPA patients and corrected for atrophy at the voxel level, we found that activation in the left dorsal frontal and parietal region and peaking in the premotor cortex was significantly (p<0.01, 5% FDR) increased in nfvPPA patients 50-150 ms after pitch perturbation onset (Figure 3B, Table 2).

**Figure 3:**
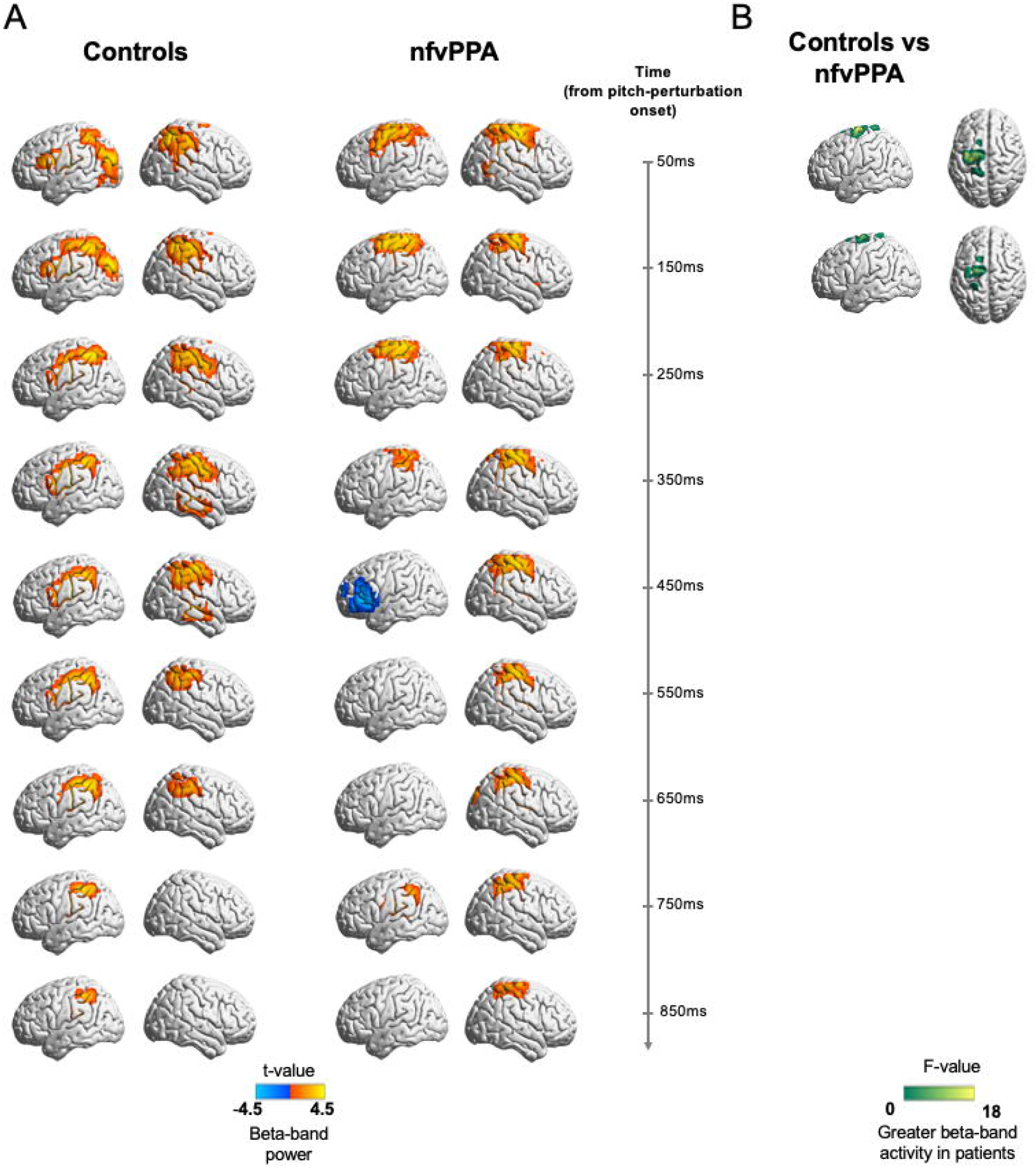
Neural activity during pitch feedback perturbation in beta band (13 - 30 Hz) **(A)** Neural activity in controls and patients with nfvPPA in the beta band (13-30 Hz) locked to pitch perturbation onset **(B)** Patients with nfvPPA (n = 17) show greater beta-band activity in the dorsal sensorimotor and premotor cortices as compared to controls (n = 14). Neural activity shown was corrected for grey-matter atrophy and was False Discovery Rate (FDR)-corrected for multiple frequency bands and time points. Cluster correction was also performed at a threshold of 20 voxels and p < 0.01. Timepoint 0 corresponds to perturbation onset.

### Neural correlates of speech motor impairment during compensation for pitch perturbation in patients

Next, we examined the associations between the abnormal neural indices of sensorimotor processing identified in the above analyses and the clinical measures of speech motor function in patients with nfvPPA. To this end we used a generalised linear model (GLM) and examined the associations between the composite speech motor score derived from clinical testing and the regional neural activity within regions-of-interest (ROI) as defined by the group contrast analyses above. The GLM between the composite speech motor score and the average peak alpha power within the right temporo-parietal junction ROI, across the timepoints where patients showed significantly reduced alpha-band activity showed a significant positive association, indicating that patients with nfvPPA with lower alpha activity are poor performers in speech motor functional tests (Figure 4A; β = 3.41, *F* = 8.31, *p* = 0.0128). A similar GLM for the associations between speech motor composite scores and the average peak beta power in the left dorsal parietal ROI, across the timepoints where patients showed significantly increased beta-band activity did not reveal a significant relationship (Figure 4B; β = -1.75, *F* = 1.72, *p* = 0.2123).

**Figure 4:**
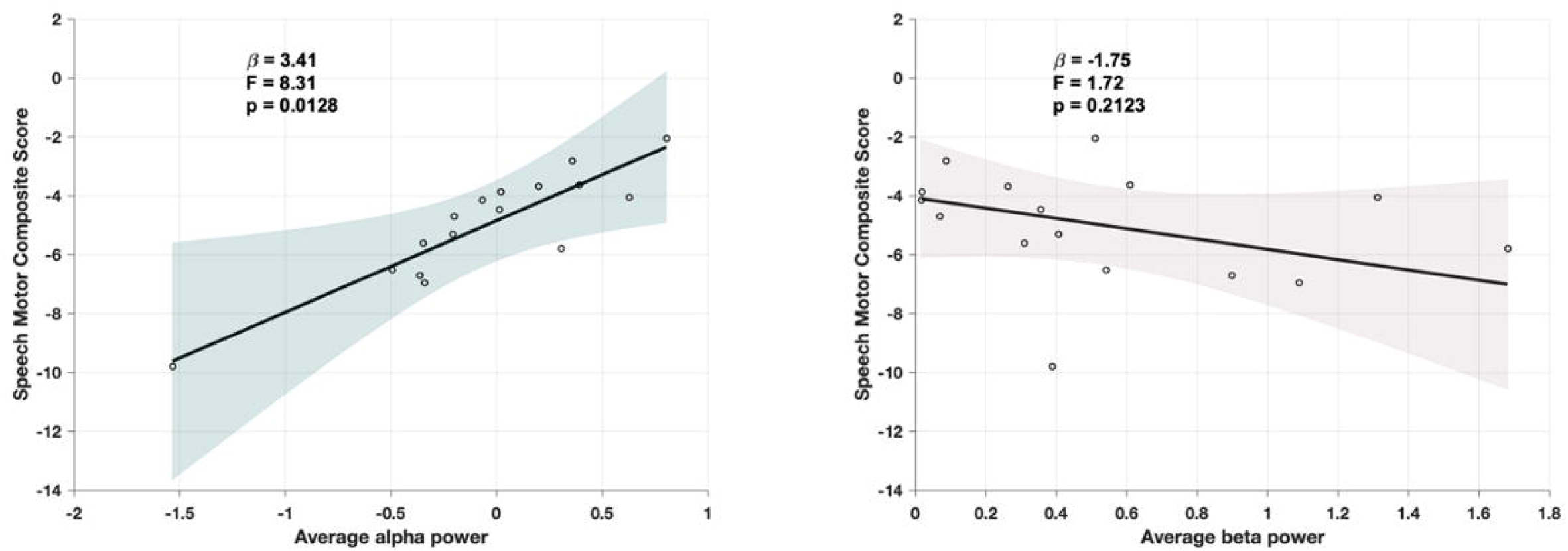
Alpha-band power predicts speech motor impairment. In a generalised linear model, patients’ speech motor composite score was significantly positively correlated with the average of peak alpha power (from 250-750ms after pitch perturbation onset) but there was no relationship with the average of peak beta power (50-150ms after pitch perturbation onset). Thus implying that the lower the induced alpha power changes due to pitch perturbation in a patient with nfvPPA, the greater is their speech motor impairment.

## Discussion

Using a novel structure-function neuroimaging approach, we investigated the cortical dynamics of speech sensorimotor control in patients with nfvPPA by examining behavioural and neural correlates of the widely-studied pitch perturbation reflex. Patients showed significantly reduced vocal behaviour compensation for altered feedback as compared to controls (only ∼33% of the peak compensation seen in controls). When we used MEG to study the neurophysiology underlying these behavioural patterns, we revealed that patients with nfvPPA had significantly reduced alpha-band activity over the right posterior temporal lobe and right temporoparietal junction that cannot be explained by cortical atrophy alone. In the beta band, patients showed significantly increased activation over dorsal sensorimotor and premotor cortices during early responses to pitch perturbation, a pattern also unrelated to cortical atrophy. Reduced alpha-band activity in patients predicted speech motor impairment in patients whereas increased beta-band activity did not correlate with speech motor impairment in patients. Together, the results suggest significant impairments in the sensorimotor integration process during speech production in patients with nfvPPA. These sensorimotor impairments may be playing a contributory role in the speech motor deficits and other characteristic loss of speech abilities associated with nfvPPA.

### Reduced vocal response to pitch perturbation

Patients with nvfPPA primarily exhibit effortful, non-fluent speech^3^ and motor speech deficits^39^ like apraxia of speech^40^ and dysarthria^41^. These deficits along with atrophy in speech motor cortices^5^ and impaired structural connectivity between brain regions involved in speech production^6-8^ suggest that the functional recruitment of the speech motor control network is impacted in nfvPPA. Models of speech motor control posit that speech production involves continuous monitoring of sensory feedback and compensation for sensory feedback prediction errors to regulate speech motor behaviour.^9, 10, 42-44^ This sensorimotor processing of the speech motor control network can be tested by altering sensory feedback during speech production and evaluating the network’s ability to compensate for this alteration. Our findings demonstrated that patients with nfvPPA have a significantly reduced vocal response to pitch feedback alteration compared to controls. This is in sharp contrast to the increased vocal response to feedback alteration shown in other neurodegenerative disorders including Alzheimer’s disease,^19, 45^ Parkinson’s disease^18, 46-48^ and cerebellar degeneration^17, 49^, when compared to healthy controls. The reduced vocal response in nfvPPA patients cannot be attributed to a limited vocal output range because patients’ within-trial and across-trials baseline pitch variability (Supp. Fig. 1) is large enough to achieve compensation. Reduced compensatory response to pitch perturbation in patients with nfvPPA may thus be an indicator of impaired sensory feedback processing and provide a useful tool to quantify such impairment.

When understanding what might be responsible for a reduced compensatory response in nfvPPA patients, it is critical to look at what triggers a compensatory response to altered auditory feedback. When auditory feedback is externally perturbed, it causes a mismatch between the predicted feedback and the actual feedback, thus generating a feedback prediction error. Any difficulty in the detection of or motor response to this error may explain a weak compensatory response to external feedback perturbation, which motivated our analyses in the neuroimaging data.

### Reduced alpha-band neural activity in patients’ posterior temporal and temporo-parietal regions and prediction of speech motor impairment

Previous studies have shown that posterior temporal regions are highly sensitive to pitch perturbation.^50-52^ It is postulated that these regions are involved in auditory error detection.^53-55^ Moreover, activity in the alpha band is thought to be associated with suppression of irrelevant sensory information^56^ suggesting alpha-modulated noise suppression in speech processing. Higher alpha-band activity indicates successful selective inhibition while less alpha-band activity would be indicative of an inability to tune out irrelevant information. Patients show significantly lesser atrophy-independent alpha-band activity than controls in the posterior right temporal lobe and the right temporo-parieto-occipital junction from 250ms (when patient’s vocal response deviates from that of controls) to 750ms, suggesting that impaired suppression of information that is irrelevant to the task at hand may be a potential mechanism contributing to their reduced vocal response to pitch perturbation. Given that the impaired reduction in alpha activity started closer to the perturbation onset and persisted until after the perturbation offset, it is likely that this abnormal neural activity hinders the processing of the auditory feedback error. If patients with nfvPPA are unable to detect the feedback error brought on by the altered pitch feedback they would subsequently be unable to integrate the error into their corrective motor output.

We also found that reduced alpha-band power over the right posterior temporal region is significantly correlated with the clinical measures of speech motor impairment in patients with nfvPPA (Figure 4). Patients who showed more hypoactivity in the cluster in the posterior right temporal lobe showed poorer scores in a speech motor composite measure as determined by neuropsychological tests.

### Increased beta-band neural activity in patients’ left dorsal sensorimotor and premotor regions

At timepoints immediately succeeding perturbation onset (50ms and 150ms), we observed an increase in beta-band neural activity in the left dorsal sensorimotor cortex and the left dorsal premotor cortex unrelated to cortical atrophy. It can be seen that patients’ vocal output at these timepoints aligns with that in controls. Thus, we believe that this increase in activity in patients did not contribute towards the reduced vocal response in patients. Although GM values in these regions did not statistically contribute to group differences in the beta band, patients with nfvPPA generally exhibit cortical atrophy in the ventral regions of the motor cortex or the laryngeal motor cortex (see Supp. Fig. 2). The dorsal hyperactivity we observe in our study may be the manifestation of an increased effort by patients in motor planning to respond to an auditory feedback error, perhaps a compensatory strategy to overcome the atrophy in regions that are normally active during laryngeal control. Indeed it has recently been shown that the conventional idea of neuronal organisation as per the ‘motor homunculus’ may not be rigid and that neurons in the dorsal motor cortex may be recruited for speech production.^57^ Moreover, the beta-band hyperactivity did not predict speech motor impairment in patients (Figure 4) which further suggests that it did not contribute to abnormal sensorimotor integration. Therefore, while differences in alpha-band activity point to a disruption in sensorimotor integration, differences in beta-band activity may be a signature of neural plasticity due to significant damage to the cortex and white matter tracts observed in nfvPPA.

## Conclusion

Here we combined MRI volumetrics with magnetoencephalographic neuroimaging and detailed behavioural analysis to identify sensorimotor control differences during pitch perturbation in nfvPPA and matched controls. Although the mechanisms connecting motor speech production abnormalities as captured in neuropsychological testing and sensorimotor integration of auditory feedback response are yet to be determined, our results suggest that sensorimotor integration abnormalities over specific regions of the speech and language network contribute to the behavioural motor speech impairments found in patients with nfvPPA. As we integrate cortical atrophy into our statistical model, these differences in alpha- and beta-band oscillatory pattern cannot be explained by neurodegeneration alone. As these findings highlight the potential for multimodal studies of structure and function, future investigations with longitudinal assessments will determine the temporal evolution and interdependencies of these sensorimotor subcomponents of speech dysfunction in nfvPPA and their progression. This knowledge will help further elucidate the mechanisms behind speech motor control and how they are affected by pathophysiological processes of neurodegeneration in primary progressive aphasia variants.

## Abbreviations

MEG: Magnetoencephalography
nfvPPA: non-fluent variant of Primary Progressive Aphasia
PPA: Primary Progressive Aphasia

## Acknowledgements

We thank the participants of this study and their caregivers for their valuable time and support.

