## Supplementary Table 1 for "Cortical dynamics of speech feedback control in non-fluent Primary Progressive Aphasia"

**Supplementary Table 1: Neuropsychological test performance in patients**

| **Test** | **Score for patients with nfvPPA*** | **Normative Score^** |
| --- | --- | --- |
| Western Aphasia Battery – Fluency (out of 10) | 6.6 ± 2.3 | 10.0 ± 0.0 |
| Western Aphasia Battery – Repetition (out of 100) | 88.3 ± 10.2 | 99.5 ± 0.9 |
| Western Aphasia Battery – Sequential Command (out of 80) | 74.3 ± 8.3 | 80.0 ± 0.0 |
| Apraxia of Speech rating (out of 7) | 3.1 ± 1.5 | --- |
| Dysarthria rating (out of 7) | 2.0 ± 1.9 | --- |
| Long Syntax Comprehension (Percentage) | 85.3 ± 18.1 | --- |
| Short Syntax Comprehension (Percentage) | 91.3 ± 18.5 | --- |
| Boston Naming Test (out of 15) | 13.0 ± 3.2 | 14.5 ± 0.7 |
| Category Fluency (Animals named / minute) | 12.9 ± 7.2 | 23.8 ± 4.3 |
| Peabody Picture Vocabulary Test (out of 16) | 14.5 ± 2.2 | 15.7 ± 0.7 |

All scores are mean ± standard deviation

*n = 19 for all scores except for Long Syntax Comprehension (n = 17) and Short Syntax Comprehension (n = 18).

^ Normative scores are from Wilson, Stephen M., et al. "Neural correlates of syntactic processing in the nonfluent variant of primary progressive aphasia." Journal of Neuroscience 30.50 (2010): 16845-16854.

For Apraxia of Speech rating and Dysarthria rating, higher scores indicate greater impairment. For all other scores, lower scores indicate greater impairment.
