## Supplementary figures and images for "Cortical dynamics of speech feedback control in non-fluent Primary Progressive Aphasia"

### Supplementary Figure 1

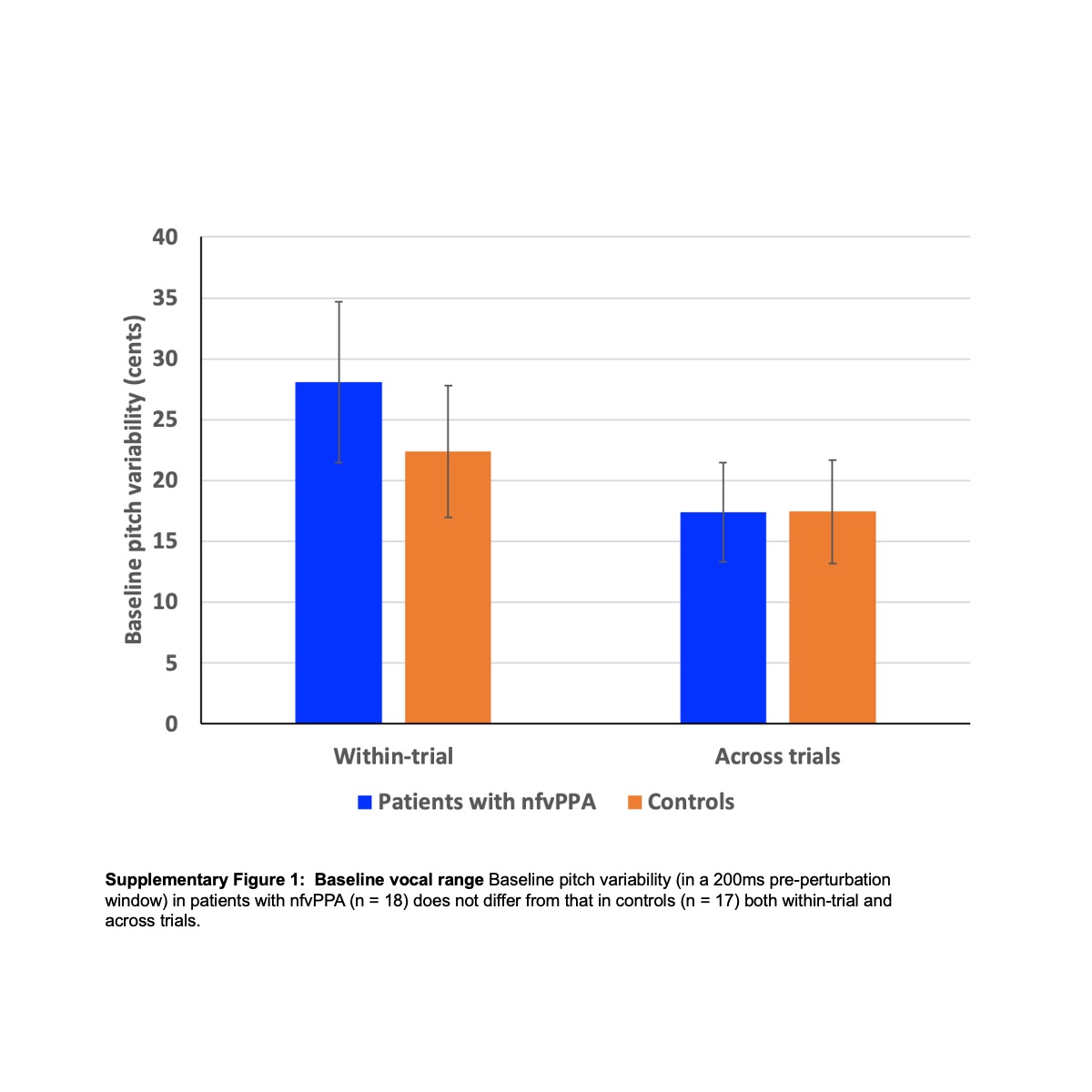

### Supplementary Figure 2

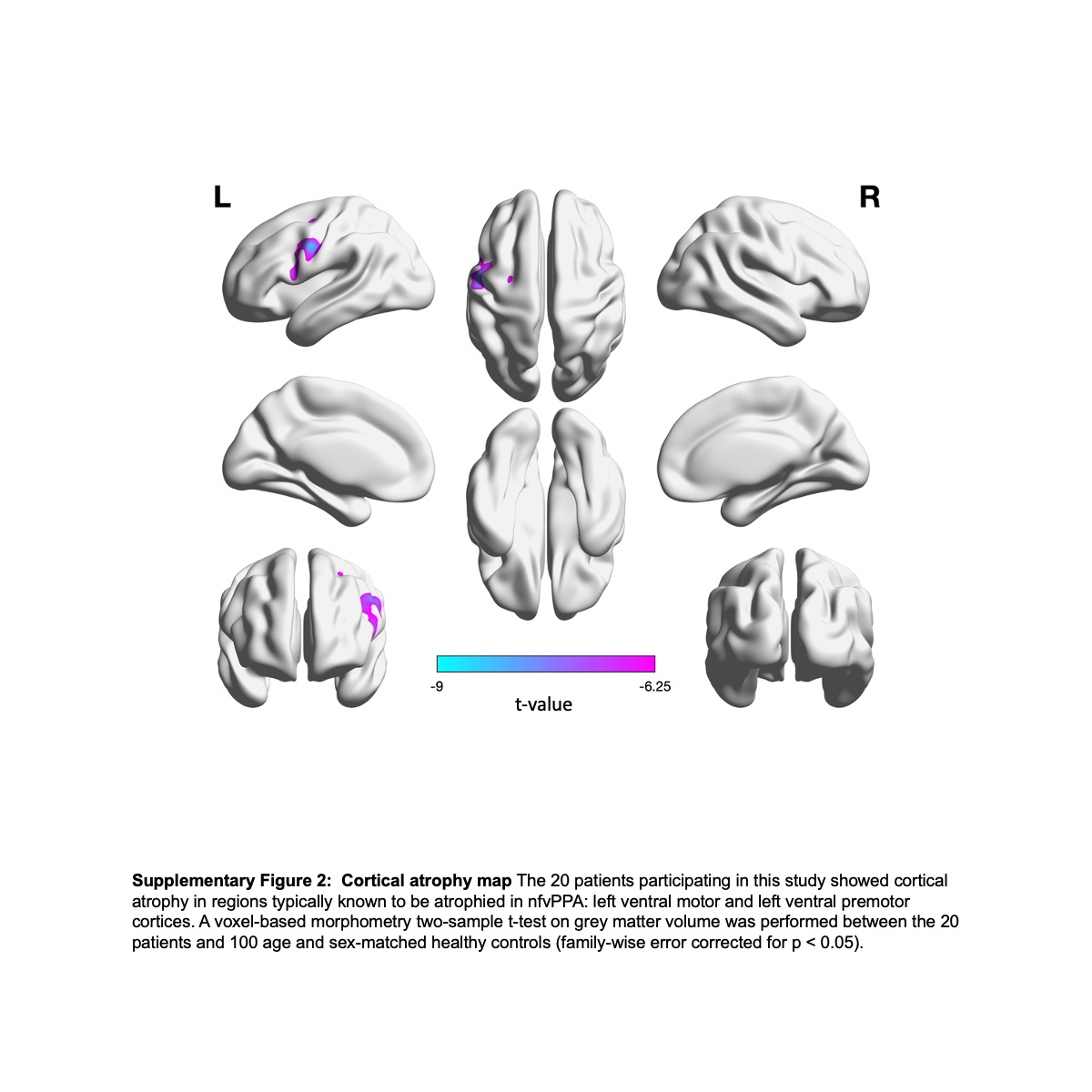
